# Maize Phyllosphere Microbial Community Niche Development Across Stages of Host Leaf Growth

**DOI:** 10.1101/161158

**Authors:** Heather C. Manching, Kara Carlson, Sean Kosowsky, C. Tyler Smitherman, Ann E. Stapleton

**Affiliations:** Department of Biology and Marine Biology, University of North Carolina Wilmington, Wilmington, NC, USA

**Keywords:** epiphytes, plant host, community assembly, Zea mays L., ecological niche

## Abstract

The phyllosphere hosts a variety of microorganisms, including bacteria, which can play a positive role in the success of the host plant. Bacterial communities in the phylloplane are influenced by both biotic and abiotic factors, including host plant surface topography and chemistry, which change in concert with microbial communities as the plant leaves develop and age. We examined how *Zea mays* leaf microbial community structure changed with plant age. Ribosomal spacer length (ARISA) and scanning electron microscopic (size trait) imaging strategies were used to assess microbial community composition across maize plant ages, using a novel staggered experimental design. Significant changes in community composition were observed for both molecular and imaging analyses, and the two analysis methods provided complementary information about bacterial community structure within each leaf developmental stage. Both taxonomic and cell-size trait patterns provided evidence for niche-based contributions to microbial community development on leaves.

## Introduction

The phyllosphere, the microbial community inhabiting the surface of plant leaves [1], can play important positive roles for the plant partner. For example, disease resistance is increased when supportive microbial bacteria are present [2,3] and increased host plant growth in the presence of specific community members has been documented [4–6], along with a positive effect of overall increased microbial diversity [7]. Leaf habitats are also extremely important for the microbial community, as this habitat is quite large in surface area globally. In temperate regions seasonal growth and annual agricultural plantings lead to repeated opportunities for colonization by adapted microbes [8,9].

Determinants of community assembly/succession in microbial systems include biotic and abiotic factors, often classified into different community assembly rules [10]. Typically random or neutral stochastic models are tested against abiotic effect models, which are called environmental, or, more recently, habitat filtering rules. Environmental filtering can produce different patterns depending on scale [11]. As recommended by Kraft et al. (2015) we will refer to environmental filtering as habitat filtering when there is no separate test for abiotic and biotic effects. A second class of assembly rules addresses the interactions of organisms, by either facilitation or competitive exclusion. These niche interactions are often modeled separately from abiotic effects, or abiotic effects are assumed to happen first.

Community assembly and succession processes are typically inferred from diversity patterns in community samples [12]. The number of different taxa (alpha-diversity), the difference in the composition of taxa between replicates (beta-diversity) and the proportion of co-occurring taxa all provide evidence for these different community-generating mechanisms. Niche construction is inferred from a pattern of decreasing beta-diversity, increasing co-occurrence, and increasing richness, while habitat filtering is characterized by decreasing beta-diversity with no trends in co-occurrence or richness [13] [14]. In contrast, neutral community assembly would exhibit increased richness (provided immigration rates are maintained), with unchanged beta-diversity and co-occurrence levels. Once we have measurements of the various components of diversity, we can compare observed trends to these expectations. The combination of increasing/decreasing trends would then support or fail to support a particular mechanism of community succession. Both trait-based (morphological, in our study) and taxonomic measures of diversity can be compared to the expected trends [15,16]. For microbial systems, cell size is one of the few traits that is measurable at the microscopic scale; other functional traits, such as metabolic activity, are more often examined at the molecular scale, by whole-community DNA sequencing, proteome analysis, or metabolite profiling.

Previous analyses of controlled inoculation phyllosphere community development indicate that habitat filtering is important, with niche effects less often detected across time, though all previous studies have used seasonal or through-time designs that confound exposure time and plant development. Pre-colonization reduces immigrant success [9,17], suggesting biotic competition is key, though high beta-diversity in replicate samples from both pre-colonized and newly inoculated communities suggested that there are many options for species that arrive first to succeed. Several crop species have been tested for seasonal patterns of phyllosphere community development. Microbial populations increased more rapidly on younger leaves of lettuce [18], and alpha-diversity increased over the first few weeks then plateaued [19], which provides evidence that habitat filtering was a contributor to community succession later in the season. In apple flowers, species richness increased early in the season and beta-diversity dropped, then rose [20]. About half the apple flower microbial species had significant co-occurrence relationships and the numbers of co-occurrences increased and decreased over time. This pattern supports niche construction as a key mechanism for apple flower colonization. As in apple flowers, across-season dicot crop phyllosphere samples became less diverse (alpha-diversity went up then declined). In the crop samples, more shared and plant-specific taxa were found later in the season [21], again supporting development of specific community structure. In contrast, Arabidopsis epiphyte beta-diversity increased with time [22], providing stronger support for stochastic effects in this system.

The history of annual opportunities for new colonization is reflected in the processes for dispersal and succession on plant leaves. Colonization of new growth in the presence of older leaves may proceed differently than colonization from distant source materials. The contributions of abiotic changes and host leaf age are confounded when sampling occurs through time, as in seasonal sampling, thus necessitating alternative experimental designs and/or additional phylogenetic or functional trait information to examine the roles of these assembly processes. Sampling serially across time, as all prior studies have done, could confound short and long-term trends and does not enable differences in plant host to be distinguished from inoculation processes [19,20]. Therefore, for this study host plant seeds were planted at different times and sampled once.

How can leaf microbial communities be measured? Molecular methods for detection of species using ribosomal fragments or sequences have become increasingly common [20,21,23]. Microscopy can be used to detect bacteria (and fungi) on corn leaf surfaces [24], and surface bacteria can be followed with reporter genes [25] as well as by direct enumeration. Microscopic or particle-based enumeration methods are especially useful for microbial communities where we have limited information on traits and horizontal gene transfer is likely [26]. For this study we used both a molecular approach to measure relatedness-based (taxonomic) microbial community composition, and direct enumeration of scanning electron micrograph (SEM) images to measure cell-size trait changes.

Our specific questions concerned the pattern of community development – addressing how community succession varies with plant age, and whether the microbial community primarily shaped by microbe-host interactions (environmental or more generally, habitat filtering) or by microbe-microbe interactions (niche construction). We found complex patterns in maize leaf microbial community structure even across the short series we sampled, with niche construction and stage-specific increasing-decreasing diversity trends observed.

## Materials and Methods

### Experimental design and sample collection

*Zea mays* B73 seeds were planted using a hand held planter between April 26^th^ and June 15^th^ 2009 at the Central Crops Research Station in Clayton, NC (http://www.ncagr.gov/research/ccrs.htm, Latitude: 35.66979° Longitude: -78.4926°). All necessary permits were obtained for the described study, which complied with all relevant regulations; permits were managed by the research station staff. Approximately twenty seeds were planted each date, with plantings between four and eleven days apart in a single field. All plants within a developmental stage were planted in the same row, bordered on either side by older and younger stages, then by additional crops, grass borders, and tree-shrub hedgerows.

All ten replicate leaf samples were collected at once on July 15, 2009. Both scanning electron microscope (SEM) and ribotype (ARISA) samples were obtained from tips of separate central leaves from ten plants at each developmental stage. SEM samples were approximately five centimeters square and were cut along the leaf mid-rib near the tip (Fig. 1). Sections were immediately placed in a solution of 2.5% glutaraldehyde in 0.1 M Sorenson’s phosphate buffer at a pH of 7.1.

**Fig. 1.**
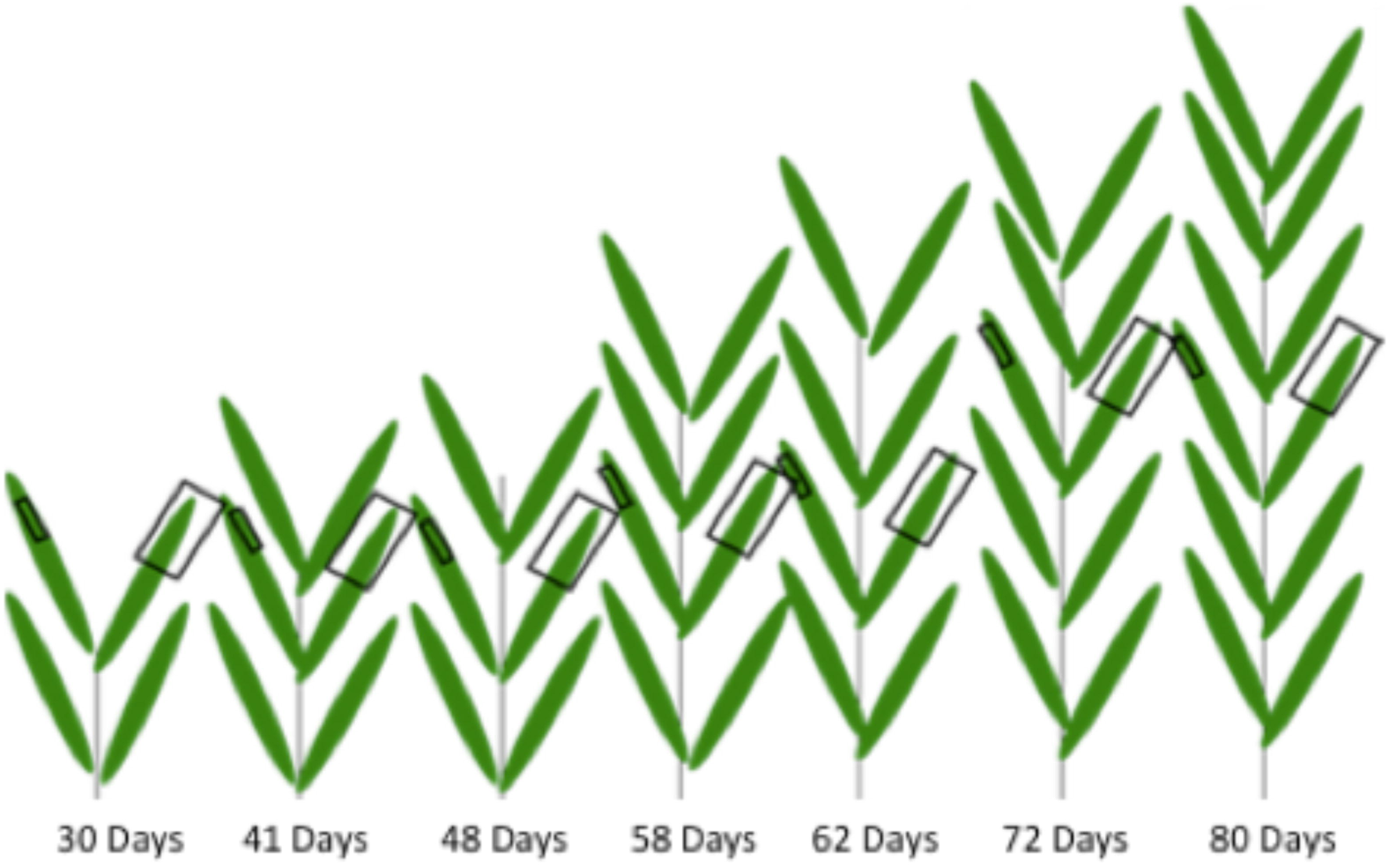
Conceptual diagram of leaf sampling procedure. Each schematized plant represents 10 plants from one of seven developmental stages. Small squares represent SEM sampling areas and large squares represent ARISA sampling areas. All samples were taken from center leaves within each developmental group and both samples per plant were taken from leaves of comparable position based on height.

Leaf wash samples for ribosomal DNA analysis were then collected from the same ten plants, using adjacent leaf numbers and a similar portion of the leaf tip (Fig. 1). Approximately 10 cm of leaf was cut, submerged in buffer (0.2mM Tris pH 7.5, 0.02 mM EDTA, 0.00012X Triton X100), agitated vigorously and removed using the method described in [27]. Leaf washes were stored on dry ice while in the field then placed in a - 70 °C freezer for long-term storage. One mock control (where no leaves were placed in the tubes) and one wash control (SEM sample from washed leaf to check for washing effectiveness) was taken for each stage.

### SEM sample processing

Excess glutaraldehyde was removed from leaf SEM samples by rinsing for 30 minutes in 0.1 M Sorenson’s phosphate buffer followed by an additional 30 minutes in distilled water. Sections were then dehydrated in an acetone series (50%, 70%, 90%, 100%), each lasting 30 minutes, followed by a 30-minute treatment in hexamethyldisilazane (HMDS). Sections were then allowed to air dry before being attached by the lower leaf surface to stubs using double-sided sticky tape. One leaf section chosen arbitrarily from each plant was sputter coated with 10 nm of platinum/palladium and observed in the SEM at 5 kV with a spot size of 3. The SEM procedure was tested prior to sampling using young (approximately 25 cm tall), greenhouse grown maize plants and *Pseudomonas aureginosa* [28].

### SEM image acquisition

Using the SEM stage control, five evenly dispersed random images (quadrants) per plant were taken at 1000x and 2000x for 35 sections (35 plants) each to give two viewpoints by which to count bacteria. All images are available from the image repository BisQue and the iPlant/CyVerse Data Commons, at doi:10.7946/P2WC77.

### Ribosomal spacer amplification and separation (ARISA)

For the DNA analysis, leaf washes were syringe filtered using previously developed methods [29] (videos of the method are available upon request) and microbial DNA was extracted using an UltraClean Microbial kit according to the manufacturer’s instructions (MoBio Laboratories, Carlsbad, CA). An equal volume of each DNA sample was amplified to ensure fair representation of the epiphytes collected.

Two separate, nested PCR reactions were performed on experimental samples as described by Cardinale et al. [30], along with four positive controls from a mixture of DNAs from cultured bacteria [31] and four negative (nuclease free water) controls. For the initial PCR step, a master mix was made to include: Q5 master mix (New England Biolabs), ITSF primer (5’-GTCGTAACAAGGTAGCCGTA-3’), LD primer (5’-CCGGGTTTCCCCATTCGG-3’), and nuclease free ultrapure water. Ten μL of master mix was added to sterile PCR tubes followed by experimental and sample controls. The samples and controls were then run on an iCycler (BioRad) PCR machine with the following settings: 1X 95°C for 30 seconds, 35X (95°C for 30 seconds; 61.5°C for 30 seconds; 72°C for 1 minute, 30 seconds), 1X 72°C for 10 minutes. For the second PCR step, a second master mix was made with the following: Q5 master mix, ITSreub primer (5’-GCCAAGGCATCCACC-3’), which contains a fluorescent HEX marker for fragment analysis in the Genetic Analyzer, SD primer (5’-TGCGGCTGGATCCCCTCCTT-3’), and nuclease free water.

Samples were run on the Applied Biosystems Genetic Analyzer and analysis of PCR product length was done using the manufacturer’s software. For each sample, two μL of PCR product and 13 μL of GS500-Rox plus HiDi (ABI, Inc, Carlsbad, CA) were combined. The samples were denatured at 94°C for ten minutes, placed on ice for two minutes, then placed into the analyzer and the program started.

### SEM image analysis

For each SEM quadrant image, Adobe Photoshop was used to manually outline each bacterial organism present (examples are shown in Fig. 2). All single bacteria present in the image were outlined; bacteria were identified by size and smoothness of circumference, as originally described by Davis (1979). Bacterial areas were measured using Image Pro Plus (Media Cybernetics, Inc., Rockville, MD, USA) and recorded for each quadrant picture that was taken. Image Pro Plus sized bacteria that were touching other bacteria as one large organism, therefore as many bacteria as possible were outlined in the clusters without outlining touching bacteria. Each image was individually measured using calibration settings based on the magnification used to take the image. Each image analyst completed analysis of a training image set and the coefficient of variation of their analyses was checked to ensure that it was less than 5%.

**Fig. 2.**
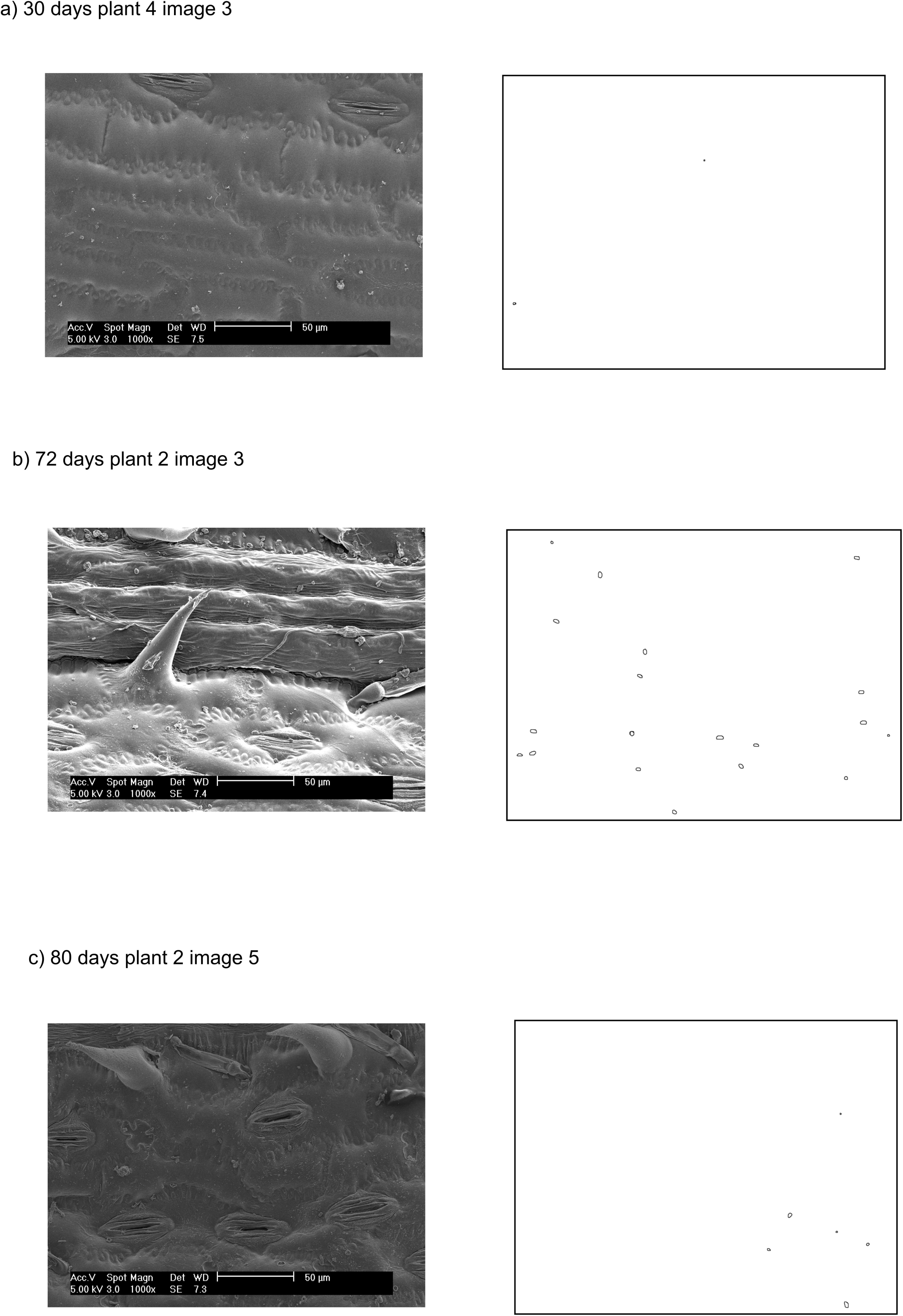
Example SEM (left panel) and bacterial overlay images (right panel). a) typical image after 30 days of growth in the field b) 72-day image, with leaf hair and stomata clearly visible c) typical image after 80 days of growth in the field.

Bacterial sizes acquired from all images were binned using methods described by Shimazaki et al. [32]. Bin sizes were assigned to the microbial samples as size-class units (SCU) and a microbial community profile was created for the samples using the RiboSort package in R [33] (R Development Core Team, 2012).

### ARISA data processing

Files from the Genetic Analyzer were opened in Peak Scanner 2 software (http://www.appliedbiosystems.com) using Analysis Method=Sizing Standard-PP, and Size standard = GS500. Only the green (hex-primer-labeled) peaks were used to determine level of bacterial abundance. The size and heights of the green peaks were calculated using the ROX GS500 standard. The individual output files were assembled into a single microbial profile file using the RiboSort package of R [34] using the script, RiboSort(data,dataformat = “standard”,dye = “G”,output = “proportions”,zerorows = FALSE,repeats = 1,mergerepeats = “presentinall”). This program automatically bins the peaks to generate abundances; we used the default threshold of 50 fluorescence units as recommended by the program authors.

### Statistical analysis of presence-absence matrices

The OTU (operational taxonomic unit) abundance matrices output from Ribosort were analyzed by Permdisp and Permanova [35–37] using Primer-E software v6 [38]. Overall square root transformation and Bray Curtis similarity resemblance were applied to the initial abundance data within Primer-E before Permanova and Permdisp tests were run. The P-values from the program were Bonferroni-corrected. Primer-E SIMPER analysis was used to compute the number of OTU contributing to differences between stages. Primer-E functions were used to compute diversity measures and taxa numbers for each sample (S1 Dataset).

The evenness (Pielou’s J’) of OTU and SCU for each sample was computed using Primer-E v6 [38]. The values of J’ for OTU and SCU were compared across plant stages by rank-transformed analysis of means using the routines in JMP Pro v11 (SAS Inc, Cary, NC) with P-values adjusted for multiple comparisons. For the rank-transformed analysis of means the P-value threshold was set at the Bonferroni-corrected value of P<0.007. This rank-transformed analysis of means is not sensitive to heteroschedasticity and is more powerful than other non-parametric tests when the data distribution is heavy-tailed [39]. Nonparametric pairwise comparisons of evenness between stages were done using the Steel-Dwass multiple-comparison method. The distributions of SCU areas in each stage of plant growth were also computed with JMP Pro v11.

### Co-occurrence analysis

Pairwise analysis was completed using the R package Co-Occur [40]; this package allows determination of the number of OTU/SCU that occur together in the same sample day at a probability above random, along with the number of taxa that are not found together (negatively correlated taxa) above the random-pair threshold. The analysis code, input and output files are available at https://github.com/skosowsky/Cooccur. We created network visualizations of the co-occurring OTU and SCU with Cytoscape (v. 3.4) and compared networks using the Cytoscape plug-in app DyNet [41]. Intensity of node connection change across the seven stages is reflected in the intensity of the color of the DyNet-analyzed nodes.

### Statistical analysis of presence-only data

The number of OTU and SCU present within each sample was computed using Primer-E v6 (abbreviated as S, also known as Chao1). The number of OTU and SCU compared to the overall mean across plant stages was analyzed using rank-transformed analysis of means via routines in JMP Pro v11 (SAS Inc, Cary, NC), with P-values adjusted for multiple comparisons. For the rank-transformed analysis of means the P-value threshold was set at the Bonferroni-corrected value of P <0.007. This rank-transformed analysis of means is not sensitive to heteroschedasticity and is more powerful than other non-parametric tests when the data distribution is heavy-tailed. Nonparametric pairwise comparisons between stages were carried out with the Steel-Dwass multiple-comparison method within JMP.

## Results

### Presence-absence matrix comparisons

*ARISA community composition and beta-diversity* Community composition changed over time in bacterial ribosomal taxa (differences in OTU, defined as ribosomal intergenic spacer length differences), with a Permanova P-value of 0.026 (S1 Table). The Permanova pairwise results for bacterial ribosomal OTU (Table 1, S2 Table) indicate that, specifically, day 30 and day 80 were significantly different from each other in community composition. Thus we can conclude there was a difference in either the richness or arrangement of the communities’ ribotypes in the earliest and latest stages. To determine which aspect of community structure contributed to the difference detected by Permanova, we examined the dispersion as a measure of beta-diversity, i.e. the amount of range in the set of species present across replicates within each time. There was no significant change in the dispersion in the ribosomal OTU datatype across stages (Permdisp P = 0.445).

**Table 1.** Permanova and Permdisp Community Structure Comparisons

| ARISA Permanova |  |  |
| --- | --- | --- |
| Stage (day after planting) |  | Significance Group* |
| 30 |  | A |
| 41 |  | A,B |
| 48 |  | A,B |
| 58 |  | A,B |
| 62 |  | A,B |
| 72 |  | A, B |
| 80 |  | B |
| SEM Permdisp |  |  |
| Stage (day after planting) | Average centroid | Significance Group* |
| 30 | 61.789 | A,B |
| 41 | 55.974 | A |
| 48 | 64.629 | B |
| 58 | 63.854 | A,B |
| 62 | 62.28 | A,B |
| 72 | 61.929 | A,B |
| 80 | 61.892 | A,B |

\*Stages with different letters are significantly different by pairwise test with a Bonferroni multiple-test-correction threshold of $P < 0.001315$ .

Evenness is a measure of the range of OTU abundances in a sample unit; we used the Pielou’s J’ measure of sample evenness for comparison of stage evenness values. Evenness values were not normally distributed and were heavy-tailed (with kurtosis less than 3), so nonparametric tests were used for analysis of evenness differences across stages. Evenness was significantly different between stages 30 and 80 in the ARISA datatype in nonparametric pairwise tests (see S1 Results), with the evenness value lower in the latest stage (i.e. a few OTU are more abundant in the stage 80 samples, compared to a more similar set of abundances in the first samples), though overall the evenness averages are not significantly different from the mean across all sample days (S1 Results). This pairwise-significant evenness difference in important OTU abundance ranges in the earliest and latest sample dates is also visible in a SIMPER analysis of ARISA OTU, with the top contributions to the day 30 OTU more evenly spread (for day 30: OTUs with 31, 19, 11, 11, 8, 6, 4%) as compared to day 80 (with OTUs of 50, 19, 10, 9, 5%, S4 Table).

*SEM community composition and beta-diversity* In contrast to the ARISA datatype, in the SEM-based SCU data the beta-diversity significantly decreased in the day 41 samples relative to the day 48 samples (Permdisp P = 0.004, Table 1, S3 Table). This beta-diversity difference confounds Permanova tests, so we cannot further compare SEM community compositions across stages with that method. The significant SEM beta-diversity community change tells us that there is some successional process at this specific stage relative to the earlier and later samples.

Comparison of the Pielou’s J’ evenness values in the SCU datatype using nonparametric tests indicated that day 72 had significantly lower evenness of size-classes relative to the overall mean (S1 Results), with pairwise tests showing that day 72 was significantly lower in evenness than day 80 (S1 Results). Pairwise comparisons among all other days were similar and not significant.

### Presence-only comparisons

We compared the number of OTU present in each sample to the overall mean using nonparametric comparisons suitable for non-normal, heavy-tailed distributions. The number of OTU – the richness – (Fig. 3a), increases significantly in the ARISA datatype at day 62 (this mean is above the confidence interval and is thus indicated with a red dot), with earlier and later samples showing no significant difference from the overall mean richness (means indicated with green dots) and no pairwise differences in means compared to each other with nonparametric tests (S1 Results).

**Fig. 3.**
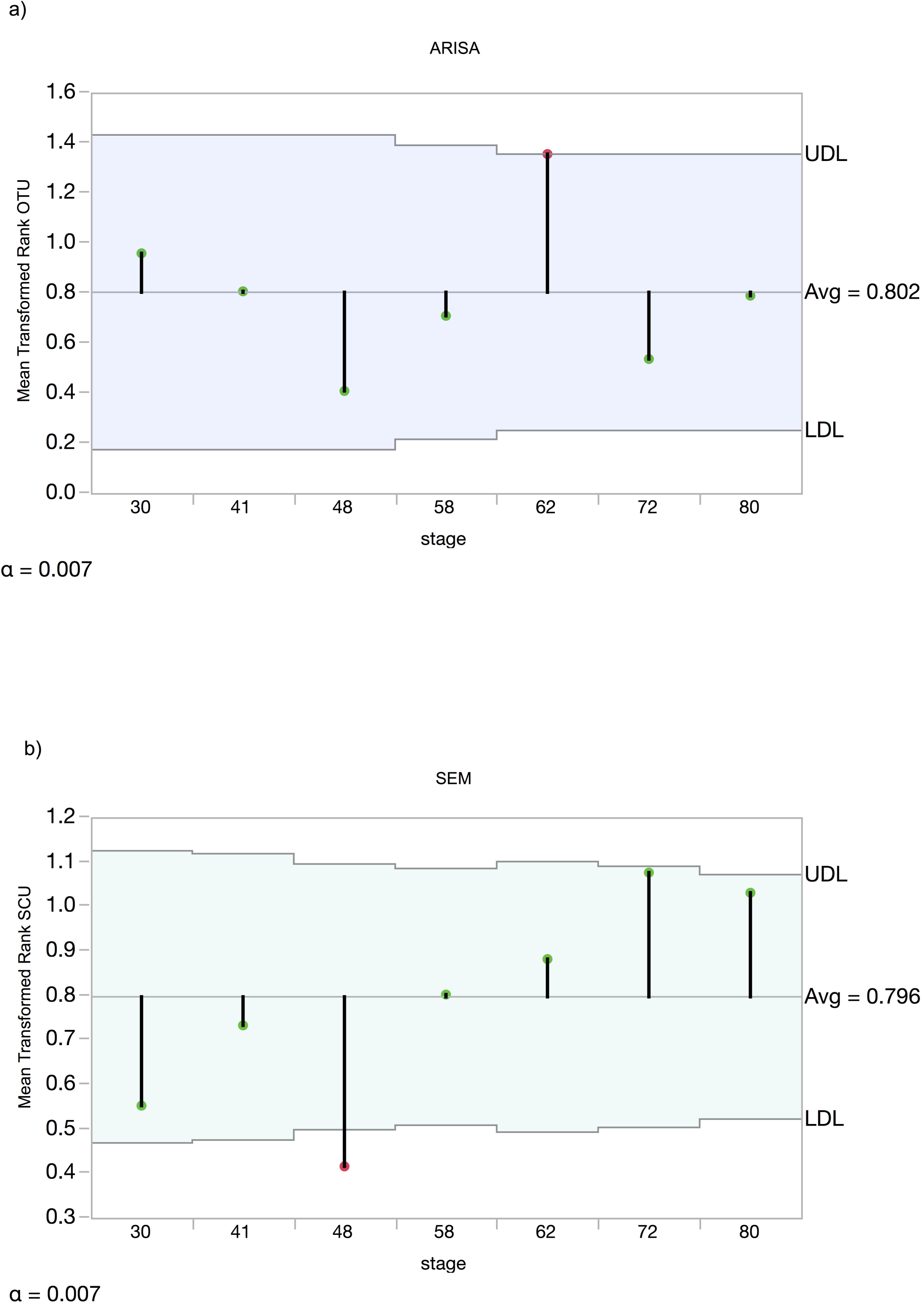
Non-parametric analysis of means of microbial OTU number (richness). Pale blue area is the confidence interval (UDL and LDL) above and below the overall mean. Rank-means for each stage are points at the tip of the drop-down bars. Significant means (adjusted P<0.05) have red line tips, while non-significant means within the confidence interval have green tips. a) ARISA mean differences with transformed ranks. b) SEM mean difference of transformed ranks of taxa.

In contrast to the ARISA datatype pattern, the number of different SCU present in the size-class datatype is significantly lower at day 48 (Fig. 3b), while all other stages are within the confidence limit for the overall mean richness. Nonparametric pairwise comparisons of SCU indicated that day 48 was significantly lower than all later-stage samples and that days 72 and 80 had significantly higher SCU means present than day 30 (S1Results).

### Core Community Members

For the ARISA datatype, there were three OTU found in all stages of community development; these were OTU9, OTU22, and OTU24, which comprised 1.5% of the total number of OTU. For the SEM datatype, there were 13 SCU (5.7% of the total); a visual display of the proportions of all 13 across the seven sample dates is shown in Supplemental Figure 3.

### Co-occurrence patterns

To further examine community assembly processes we looked at trends in the co-occurrence of pairs of species in both phyllosphere datatypes. We examined these patterns in our data to identify ways that community composition could differ. Positive co-occurrence is defined as species occurring together more often than random (and thus having potential mutualistic interactions), while negative co-occurrence values indicate that the species pair is found together less often than the random expectation (potential antagonistic interactions). Co-occurrence patterns changed across plant stages in both ARISA (Fig. 4) and SEM (Fig. 5). Participation of certain OTU and SCU in varying interactions across multiple stages is indicated by the red color intensity and number of overall edge interactions in the DyNet networks (Fig. 4b and Fig. 5b). In the ARISA datatype, highly variable co-occurring OTU (intense red color in Fig. 4b) had both positive and negative interactions with other community members. In contrast, the SEM datatype exhibited more positive interactions connecting highly dynamic nodes (intense red nodes in Fig. 5b). There were more positive than negative significant co-occurrences in both the ARISA and SEM datatypes (Fig. 4 and 5, edge arrows with points). The number of co-occurrences increased later in the season in each datatype, though not at the same stage; as seen in Fig. 4, ARISA had the largest number of network nodes at day62 and for SEM (Fig. 5) the largest number of nodes was day72. The overall proportion of co-occurring pairs was similar in each datatype (Table 2), with a minority of microbial types found to have significant co-occurrence partners. Specific OTU (S1 Figure) and SCU (S2 Figure) identifiers are individually displayed in stacked plots that show details about which OTU occurred in more than one pair.

**Table 2.** Proportion of Co-occurring Pairs within the ARISA and SEM Community Measures

|  | Total number of OTU or SCU | Number that significantly co-occur at least once | Percent with no interaction |
| --- | --- | --- | --- |
| ARISA OTU | 195 | 14 | 93% |
| SEM SCU | 229 | 28 | 88% |

**Fig. 4.**
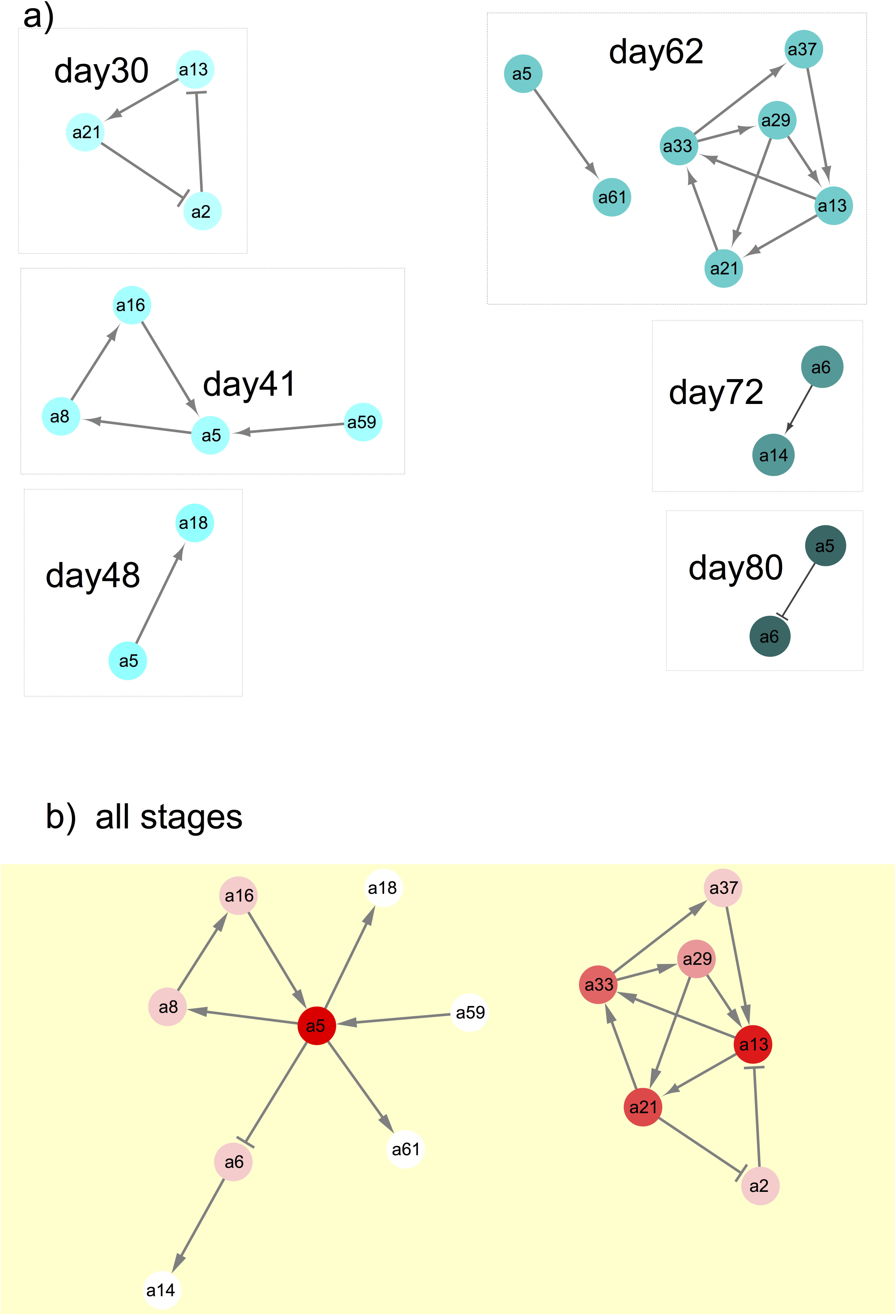
Network visualization of significantly co-occurring OTU pairs at each plant stage, with the identifier number for each OTU shown with the label “a#”. If there were no significant co-occurring OTU, then that sample stage was not shown. Negative interactions between OTU nodes are shown as edges with a barred arrow tip and positive correlations between OTU are indicated with pointed arrows. a) Significant co-occurring OTU from each sampled stage. b) Results of DyNet analysis of co-occurring OTU most changed across stages. Only significant co-occurring OTU (as shown in part a) were analyzed using DyNet. Intensity of the red color of nodes indicates amount of change across stages, with darker color indicating the most ‘rewiring’ across stages.

**Fig. 5.**
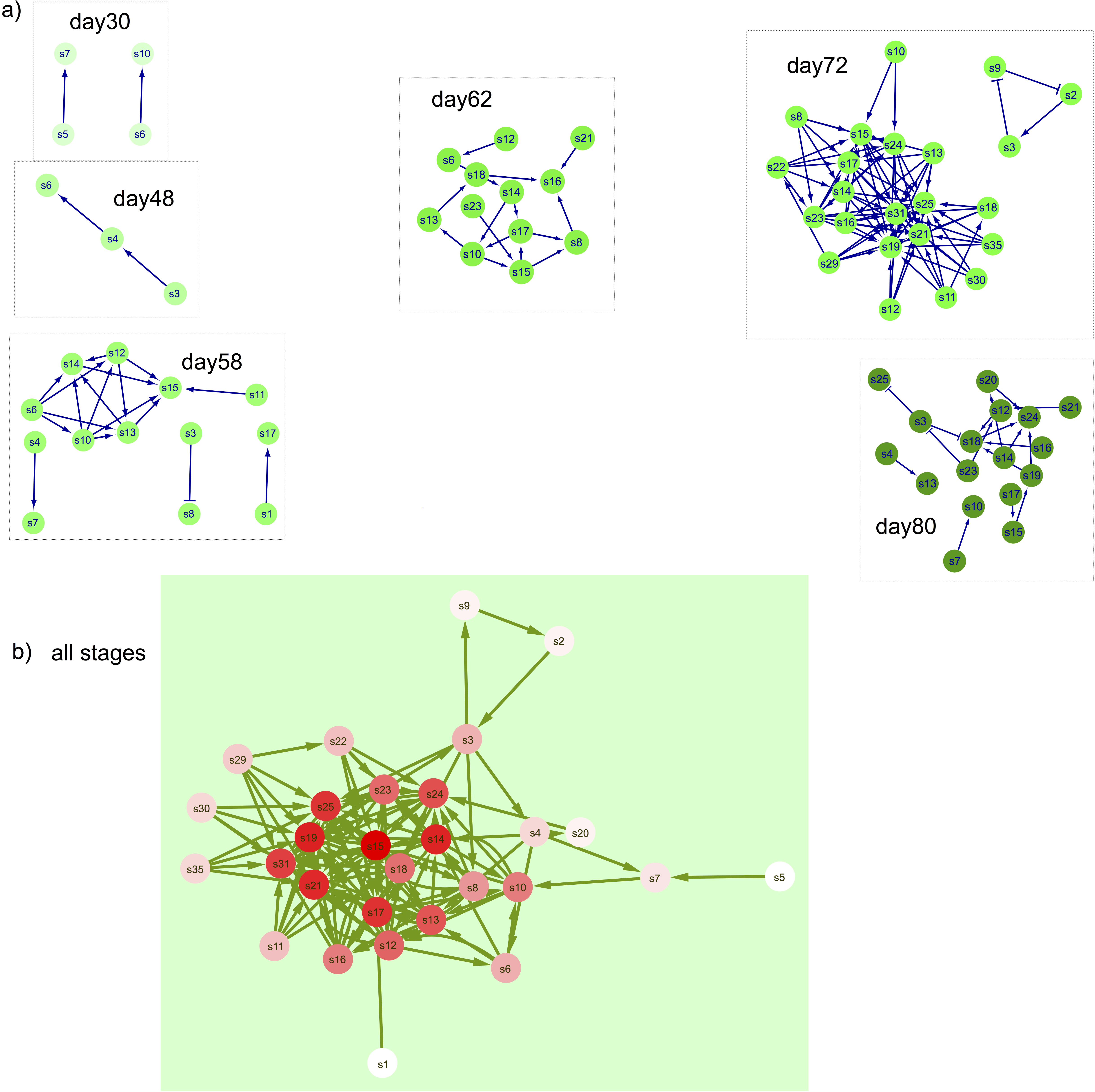
Network visualization of significantly co-occurring SCU pairs at each plant stage, with the identifier number for each SCU shown with the node label “s#”. If there were no significant co-occurring SCU, then that day was not shown. Negative interactions between SCU nodes are shown as edges with a barred arrow tip and positive correlations between SCU are indicated with pointed arrows. a) Significant co-occurring SCU from each sampled stage. b) Results of DyNet analysis of co-occurring SCU most changed across stages. Only significant co-occurring SCU (as shown in part a) were analyzed using DyNet. Intensity of the red color of nodes indicates amount of change across stages, with darker color indicating the most ‘rewiring’.

## Discussion

Our experimental design allowed for a single microbial sampling that was not confounded by differential exposure to recent weather events. This design, which has recently been recommended by Shade et al. (2013) and by Williams et al. (2013), allows us to examine the contributions to community assembly due to dispersal, filtering and niche-related processes. Comparisons of presence-absence patterns to presence-only richness within a datatype are complementary analyses, as the presence-absence matrix allows for overall tests of community difference (Permanova-based) and tests for beta-diversity difference as difference in homogeneity of variance (Permdisp). Presence-only lists of taxa or size-classes allow examination of the richness dimension of the microbial communities, abstracted away from other structural differences.

Are two datatypes better than one? We measured a trait (microbe size) and a relatedness (ribosomal length difference) metric, and the two datatypes were not especially similar at specific time points, making them complementary rather than redundant; we infer that predictive power from one datatype to the other would be low. The differences between datatypes suggest that size-classes may include different taxa, in other words our size-classes may not have a phylogenetic signal. Some recent work in phytoplankton and soil microbial communities contrasts with our results, showing that traits can be conserved across taxonomic groupings [26,42]; generally conservation of traits and evolutionary relatedness is likely to depend on the genetic architecture of the phenotypic trait [43]. The relationship between metabolic and functional traits and phylogenetic signal is as yet unclear for microbial communities that have substantial levels of gene transfer [44] [45]. Complementary information provided by two (or more) datatypes is thus recommended for future phyllosphere experiments.

Misclassification of ribosomal spacer lengths to different taxa when they are in fact due to population differences, and misclassification of large bacterial cells in a size-class separate from their smaller offspring, would have consequences for trends across samples if the misclassification probability were different at the different stages of community development. In our study, the underlying assumption is that our probability of misclassification is the same across stages. For size-classes, misclassification could increase with time due to growth of cells, resulting in a continuous increase in the number of SCU over time. We do not see this trend in our data, so from our study there is no evidence of confounding misclassification. We cannot rule out misclassification that occurs only at specific stages using our experimental design. We suggest that sample processing for future experiments include in-line size separation so that size-classes, ribosomal data, and molecular/metabolic data are collected as true multivariate data in each sample. Image information such as our scanning electron micrographs or fluorescence-based microbial images could also be analyzed for patterns that could illuminate resource use traits and specific cell-shape-based mutualisms [46].

Trends in richness, co-occurrence trends, and beta-diversity changes together define overall community assembly mechanistic contributions, with habitat filtering characterized by unchanged richness, unchanged co-occurrence frequency, and decreasing beta-diversity. Niche development would be observed as a combination of increasing richness, increasing co-occurrence, and lowered beta-diversity. Finally, a neutral pattern -- with no change in immigration probability over time -- would exhibit trends toward increasing richness, unchanged co-occurrence, and unchanged beta-diversity. Comparison of recent phyllosphere analyses and our work using this framework emphasizes the role of niche development as a mechanism of community assembly.

First we compare our results with recent work in apple flower microbial communities, where beta-diversity was initially at relatively high levels, decreased, then increased over the remainder of the season; richness (measured as phylogenetic diversity in the apple flower work) increased then leveled off, and finally co-occurrences showed early increase then decline and rebound toward higher levels over the season [20]. To summarize, early in the apple flowering season the diversity patterns support niche development mechanisms, and later in the season both habitat filtering and niche development patterns were seen. In our study of maize leaf microbial epiphytes we also found that there was no overall temporal trend. Our leaf ARISA pattern of diversity change was not especially similar to the apple flower changes; for example, we found no significant change in beta-diversity, while beta-diversity declined then rose over the later part of the season in apple flowers. Our ARISA richness and the apple flower phylogenetic diversity also differed, with apple flower phylogenetic diversity increasing then leveling off, while on maize leaves richness increased only in one later growth stage. The evenness in both our ARISA and apple flower phylogenetic diversity varied; in apple flower the ‘mid’ stage had a lower evenness while in our ARISA dataset the day 80 evenness was lower than in day 30. Co-occurrences in our study increased in mid-season, as did the apple flower communities in the early flower stage. Overall, the early season patterns in the across-season apple flower microbial community analysis were the most similar in community assembly to our maize leaf staggered-stage analysis. In addition to the apple flower study, in a recent study of three different crop leaf phyllosphere seasonal patterns, Copeland et al. [21] found that alpha-diversity increased then decreased across the season, with more shared species later in the season. In all three crops, beta-diversity (as measured as Unifrac distance for leaf-specific microbes) decreased across season, though not evenly; the beta-diversity decrease was more prominent later in season and at one time-point was higher during the later phase. Co-occurrences were not analyzed in this study, so these crop study results are consistent with niche and/or habitat mechanisms.

Core communities in crops are those that are either recruited early and retained, or continually input across the season. These are often found by presence [47], as in our analysis of ARISA and SEM core microbial community members. We found a relatively small proportion of core OTU and SCU; other phyllosphere studies such as on apple flowers [20] found more (for apple flowers the generalist category was 14%, for example). Our core set should be considered as a first description, as we did not consider absences or compare our core set to air or soil samples. Instead, our core set focused on microbial community members that colonized leaves of all developmental stages.

Our ARISA taxonomic diversity patterns and microbial cell size trait datatypes exhibited different specific patterns of diversity, but there was similarity in trend overall, as both datatypes support niche process contributions to community assembly. Niche (deterministic) processes have recently been found to be important in soil microbial communities [48,49]. In lakes, microbial generalists found at multiple spatially separated sites had a better fit to neutral stochastic models, with specialists exhibiting a more deterministic pattern [50]. As leaf surface communities are quite different than air or soil communities [8], and the number of core species in our inventory is low, we suggest that many maize leaf microbial community members are specialists. Our staggered experimental design highlighted the role of microbe-microbe interactions in leaf microbial community development. We thus suggest that future experiments use staggered experimental designs and multivariate metabolic, genic, and size-trait data collection.

## Acknowledgements

AS conceived the experiment and the data analysis and wrote the paper; HM designed and supervised the SEM image analysis; CS carried out the field experiment, extracted leaf wash DNA, and collected the SEM images; KC amplified the ribosomal fragments and analyzed the ribotype data; SK carried out the co-occurrence analyses. All authors edited the manuscript.

We are grateful to A. Ulloa-Bacillio and J. Geyer for assistance with SEM image analysis and to the staff of the Central Crops research station for their superb care of the experimental fields.

**Supporting Information Captions**

S1 Readme. List of all supporting information filenames, description of file content, column header definitions, in doc format.

S1 Dataset. Taxonomic or size-class unit diversity values for each sample.

S1 Table. ARISA Permanova results output for overall model fit.

S1 Figure. ARISA co-occurrence stacked block graphic showing each OTU pair.

S1 Results. Full statistical output tables for ARISA and SEM analyses.

S2 Figure. SEM co-occurrence stacked block graphic showing each OTU pair.

S3 Figure. SEM cell plot of SCU that occurred in all sampled days.

S2 Table. ARISA Permanova results output for all pairwise tests.

S3 Table. SEM datatype multivariate dispersion (Permdisp) test results.

S4 Table. ARISA SIMPER results listing contributions of each OTU to stages.

*Files in the CyVerse/iPlant Data Commons repository, http://mirrors.iplantcollaborative.org/browse/iplant/home/shared/data_commons/Stapleton_MaizeLeafMicrobesByLeafAge_2015*

(doi:10.7946/P2WC77):

*Supplemental_SEM_files Folder:*

*binnedSEMSizes.csv*

*readme_SEM_folder_file_metadata.txt*

*SEM_Images Folder and Subfolders*

*SEM_overlays Folder*

*SEM_ribosort_input_files Folder*

*SEM_RiboSort_output_files Folder*

*Supplemental_ARISA_files Folder:*

*ARISAfilenamemetadata.csv*

*ARISA_Ribosort_input Folder*

*ARISA_Ribosort_output Folder*

*ARISA_runs Folder and SubFolders*

*readme_ARISA_folder_and_file_metadata.txt*

